# Neural reinstatement tracks attentional selection of object features in working memory

**DOI:** 10.1101/2021.11.02.466810

**Authors:** Frida A. B. Printzlau, Nicholas E. Myers, Sanjay G. Manohar, Mark G. Stokes

**Author notes:** Corresponding author: Frida Printzlau.

## Abstract

Attention can be allocated in working memory (WM) to select and privilege relevant content. It is unclear whether attention selects individual features or whole objects in WM. Here, we used behavioural measures, eye-tracking and electroencephalography (EEG) to test the hypothesis that attention spreads between an object’s features in WM. Twenty-six participants completed a WM task that asked them to recall the angle of one of two oriented, coloured bars after a delay while EEG and eye-tracking data was collected. During the delay, an orthogonal ‘incidental task’ cued the colour of one item for a match/mismatch judgement. On congruent trials (50%), the cued item was probed for subsequent orientation recall; on incongruent trials (50%), the other memory item was probed. As predicted, selecting the colour of an object in WM brought other features of the cued object into an attended state as revealed by EEG decoding, oscillatory *α*-power, gaze bias and improved orientation recall performance. Together, the results show that attentional selection spreads between an object’s features in WM, consistent with object-based attentional selection. Analyses of neural processing at recall revealed that the selected object was automatically compared with the probe, whether it was the target for recall or not. This provides a potential mechanism for the observed benefits of non-predictive cueing in WM, where a selected item is prioritised for subsequent decision-making.

---

Working memory (WM) allows us to maintain and process information over the short-term (Baddeley, 2003). Selective attention plays a key role in top-down control of WM by biasing processing toward task-relevant information for storage and retrieval (Baddeley, 2012; Gazzaley & Nobre, 2012).

Early evidence for internal attention toward content in WM came from studies showing that retro-cueing relevant information after encoding improves memory performance (Griffin & Nobre, 2003; Landman, Spekreijse, & Lamme, 2003). There is substantial overlap between behavioural and neural mechanisms associated with attention to stimuli in the environment and internal attention toward content in WM (Chun & Johnson, 2011; Kiyonaga & Egner, 2013; Panichello & Buschman, 2021). In external visual attention, object-based theories propose that objects are a key unit of attentional selection (Desimone & Duncan, 1995; Duncan, 1984). When attention is directed toward one object-feature, neural processing is enhanced for both task-relevant and task-irrelevant features of that object, suggesting perceptual attention automatically spreads between an object’s features (Ernst, Boynton, & Jazayeri, 2013; O’Craven, Downing, & Kanwisher, 1999). Whether internal attention in working memory is similarly object-based remains a topic of debate.

A wealth of evidence indicates that individual features of objects can be prioritised in WM. Neural activity preferentially codes for task-relevant features during the WM delay, indicating that feature-based attention may enhance task-relevant features (Bocincova & Johnson, 2019; Serences, Ester, Vogel, & Awh, 2009; Woodman & Vogel, 2008). Further support comes from behavioural studies showing that participants’ memory judgements are improved if a retro-cue informs them which feature dimension will be probed after a delay (Hajonides, van Ede, Stokes, & Nobre, 2020; Niklaus, Nobre, & Van Ede, 2017). While these studies show that individual features can be prioritised in WM, it is possible that the benefit arises because valid cueing paradigms allow irrelevant features to be “dropped” or removed from memory (Souza & Oberauer, 2016). It is less established whether attentional selection spreads to other features belonging to the same object, as in object-based attention, when uncued features cannot be dropped in this way.

Studies employing non-predictive cueing have found that attentional enhancement spreads between different locations on an object (B. Peters, Kaiser, Rahm, & Bledowski, 2015) and between an object’s features in WM (Zokaei, Manohar, Husain, & Feredoes, 2014; Zokaei, Ning, Manohar, Feredoes, & Husain, 2014), consistent with object-based selection. Using an ‘incidental task’ to retro-cue a single feature of a multi-feature WM object in a non-predictive way, Zokaei et al. (2014a, 2014b) showed that subsequent memory performance improved for another feature of the same object. Such obligatory enhancement of associated object-features is predicted by theoretical models of attention in WM that rely on associative pattern completion by neural attractors (Lansner, Marklund, Sikström, & Nilsson, 2013; Manohar, Zokaei, Fallon, Vogels, & Husain, 2019), unlike models that activate features individually (Bays & Taylor, 2018). While these findings are consistent with object-based selection, behavioural paradigms alone face a challenge when it comes to assessing the relative priority status of features during the maintenance delay. For example, incidental cueing effects could also arise if the cued feature remains more accessible in memory thereby facilitating subsequent retrieval. Examining the neural effects of incidental cueing will help establish whether neural processing is enhanced for other features of the cued object, as expected if attention spreads between an object’s features in WM.

Neuroimaging studies have revealed neural markers of attentional orienting in WM following valid retro-cues. First, predictive retro-cues engage spatial attention toward the memorized position of a cued object, indexed by contralateral suppression of α-power and directional gaze bias (Myers, Walther, Wallis, Stokes, & Nobre, 2015; Poch, Campo, & Barnes, 2014; van Ede, Chekroud, & Nobre, 2019). A recent study showed that a non-predictive colour cue may similarly elicit gaze bias, though smaller in magnitude (van Ede et al., 2020), but whether non-predictive cues modulate α-lateralization remains unestablished to our knowledge. Second, WM content cued as task-relevant is represented more strongly in neural patterns as decoded from blood oxygen-level dependent and EEG activity relative to uncued content (LaRocque, Lewis-Peacock, Drysdale, Oberauer, & Postle, 2013; Lepsien & Nobre, 2007; Lewis-Peacock, Drysdale, Oberauer, & Postle, 2012). It is unknown whether selection of a single feature by a non-predictive retro-cue similarly reinstates associated features of the cued object during the WM delay, even though these are no more relevant than features of another WM object.

In the present study, we combined behavioural methods, EEG and eye-tracking to track spatial and feature-based attention during the memory delay and investigate whether attentional selection spreads between an item’s features in WM. Participants performed an orientation recall WM task with an ‘incidental’ delay-task serving as a non-predictive colour retro-cue. Importantly, the cue did not require retrieval of any other features and did not predict which item would be probed for orientation recall. On congruent trials (50%), the cued item was also probed for recall; on incongruent trials (50%), the uncued item was probed. If attention spreads between an item’s features in WM, selection of a single feature during the delay should enhance memory performance and neural processing of other features of the same object. As predicted, selecting the cued colour for the incidental task shifted other features of the cued object into an attended state. This was revealed by improved orientation recall, enhanced orientation decoding from EEG, lateralized α-power, and gaze bias relative to the cued item’s position.

In a second set of analyses, we examined how selection of the cued object during the delay affects neural processing of the probe stimulus at recall. Several recent theories propose that attention prioritises information in WM for decision-making (Myers et al., 2017; Heuer et al., 2020; Olivers & Roelfsema 2020). Sensory input elicits a stronger neural response in sensory cortex if it matches relevant information in WM (Gayet et al., 2017; Hayden & Gallant, 2013) and attended WM objects automatically guide visual attention, while “accessory” items that might become relevant later do not (Olivers, Peters, Houtkamp, & Roelfsema, 2011; J. C. Peters, Goebel, & Roelfsema, 2009). In the present study, the cued item was no more relevant to orientation recall than the uncued item, but the cued item’s selection for use in the incidental task may nevertheless place it in a special state for interacting with new sensory input (Heuer, Ohl, & Rolfs, 2020; Olivers & Roelfsema, 2020).

Analyses of the neural dynamics at recall showed that a comparison-signal was computed between the cued item and the probe stimulus, whether it was the target for recall (congruent) or not (incongruent), facilitating WM performance on congruent trials and interfering with performance on incongruent trials. These results suggest the selected object was prioritised for subsequent decision-making, providing a potential mechanism for non-predictive cueing effects in WM.

## Material and methods

The hypotheses and methods were pre-registered on the Open Science Framework after data collection had begun, but prior to data analyses (https://osf.io/pgdfj). The pre-processing pipeline was tested on a pilot participant not included in the analyses. The methods are as described in the pre-registration, unless otherwise specified in text and under ‘deviation from pre-registered methods’ section. Most notably, we planned to recruit a total of 35 participants, but due to Covid-19 restrictions only 32 participants completed the experiment.

### Power analyses

Pre-registered power calculations were based on a previous dataset that decodes the orientation of two WM items using EEG (Experiment 1; Wolff, Jochim, Akyürek, & Stokes, 2017). Re-analysis of the data using the pre-registered methods for this study revealed an expected effect size of *d*=0.43 to detect a difference in orientation decoding quality between a cued and an uncued item. Consequently, in this study, to detect an effect size of 0.43 with a power of 0.8 using a paired samples t-test (one-tailed, cued > uncued), the planned sample size was n=35. Power calculations were performed using G*Power (Faul, Erdfelder, Buchner, & Lang, 2009).

### Participants

A total of 32 participants completed the experiment (see *Deviations from pre-registered methods*, 1). One participant was excluded from analyses due to colour-blindness. A further five participants were excluded as they had less than 80% of usable epochs remaining after artefact rejection, in line with the pre-registered exclusion criteria, leaving a total of 26 participants for analyses (14 female, 12 male). Participants were aged between 18 and 34 (M=23.23, SD= 5.02). Twenty-two were right handed and four were left handed. All participants had normal or corrected-to-normal vision (including normal colour vision), and did not have any history of neurological or neuropsychiatric disorders. Participants provided informed consent prior to participating in the study. They were reimbursed for their time at a rate of £15/hour or with course credit. The study has been approved by the Oxford Central University Research Ethics Committee (R55073/RE008).

### Procedure

Participants performed a computerised experimental task (Figure 1A). It consisted of a continuous report WM task, in which participants reported the orientation angle of one of two items in memory at the end of a delay. During the delay, participants performed a secondary “incidental task”, where a non-predictive colour cue, presented centrally, required a speeded mismatch response: does the colour differ from the colours of the two memory items? At the end of the trial, a probe indicated which of the two items participants should report. This was either the same item as the one that was cued on the incidental task (congruent condition, 50% of trials) or the other item in memory (incongruent condition, 50% of trials).

**Figure 1.**
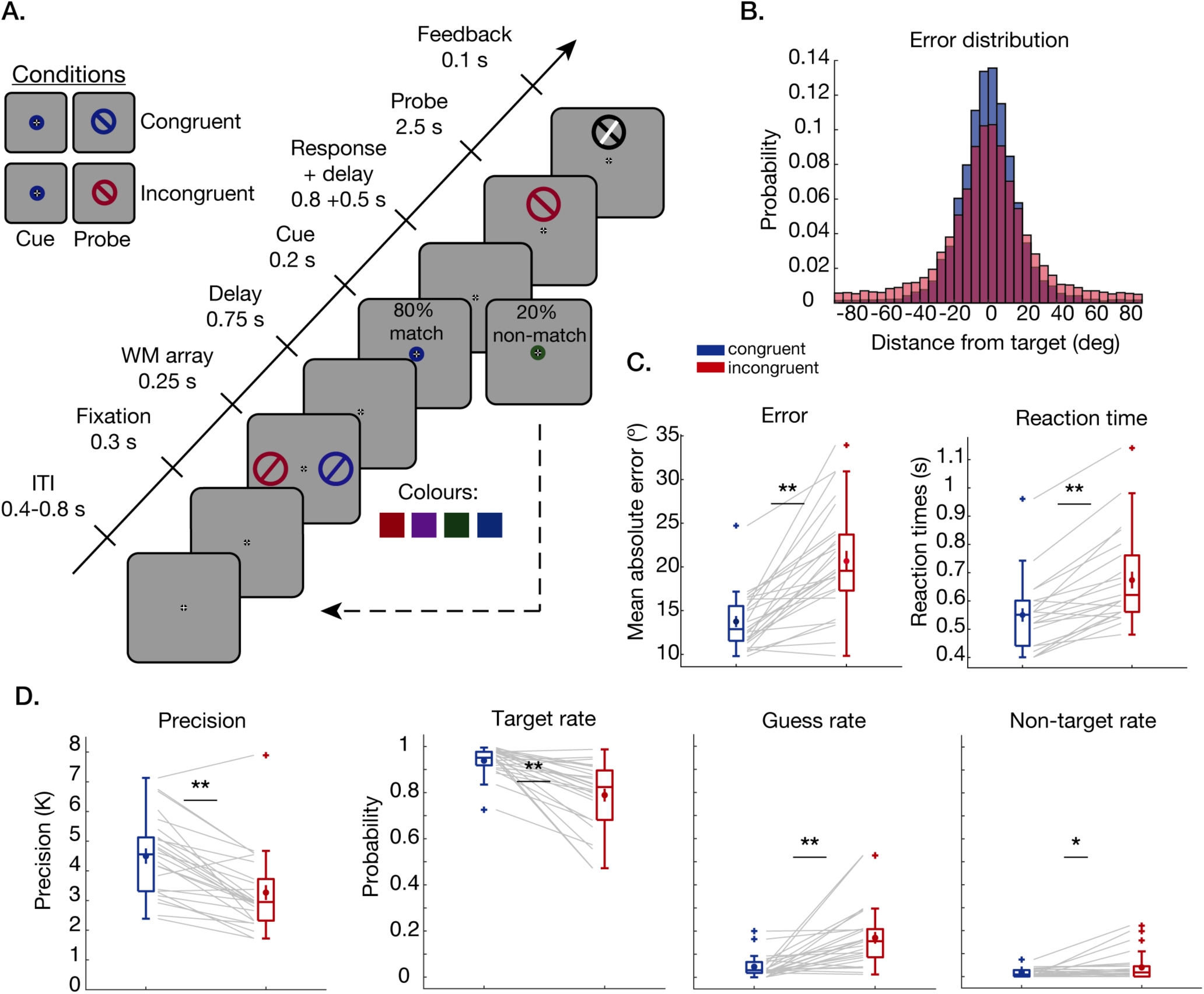
Task design and behavioural results. **A**. Task design. Participants remembered two oriented, coloured bars on each trial, presented laterally. During the maintenance delay, they were shown a non-predictive colour cue that asked them to make a speeded mismatch-response (i.e., does it differ from the colours in memory?; ‘incidental task’). On non-match trials (20%) and match trials with incorrect responses, the rest of the trial was skipped (dashed arrow). On the remainder of trials (80%) the response window was followed by another delay and a recall probe prompting participants to report the orientation of one of the items, cued by colour. This item was either the same as was also cued on the incidental task (congruent condition, 50%), or the other item (incongruent condition, 50%). **B**. Histogram of angular error (deg) relative to target orientation across all participants for congruent (blue) and incongruent (red) trials. **C**. Left panel: boxplot of mean absolute error. Right panel: boxplot of median reaction times (time until response initiation). Overlaid mean and SEM. Grey lines show individual participants. **D**. Swap model parameters. Boxplots of precision (K), target rate, guess rate and non-target rate. \**p*<.05, \*\**p*<.001.

Following an initial fixation period of 300 ms, participants were presented with a WM display consisting of two oriented, coloured bars. The WM items were presented on screen for 250 ms followed by a delay period of 750 ms. Then, for the incidental task, participants were shown a central coloured ring as a cue for 200 ms. Participants indicated whether the colour of the cue matched the colour of either of the bars in memory. On mismatch trials (20%), participants pressed the space bar within 1000 ms of incidental cue onset, after which the rest of the trial was skipped. Participants received feedback for correct responses or incorrect omissions, in the form of a happy or sad smiley face for 200 ms. On match trials (80%), participants made no response and the trial continued (unless they accidentally pressed the space bar, in which case the trial was also discontinued). After another 500 ms (1500 ms after cue onset), the probe appeared on the screen. The colour of the probe indicated which of the two items participants should report. Participants used the arrow keys to turn the on-screen dial to match the orientation of the probed item and pressed the space bar to respond. There was no maximum time to respond, but once participants initiated a response the trial would time-out within 2500 ms. They received visual feedback of the correct orientation for 100 ms. The inter-trial-interval was uniformly jittered between 400-800 ms. At the end of each block, they received feedback of their incidental colour change-detection rate and on their mean absolute error on the orientation recall task.

Participants completed a total of 1400 trials. The 20% non-match incidental trials were excluded, leaving 1120 trials per participant for analyses. Participants practiced the task beforehand until they reached a criterion of >80% correct incidental task change-detection rate and <25 degrees mean absolute error relative to the target orientation.

### Apparatus and stimuli

The task was programmed and stimuli presented in Matlab with Psychophysics Toolbox (Brainard, 1997; The MathWorks Inc., 2017). The task was presented on a 24-inch, 1920×1080 pixel monitor running at 100Hz.

Stimuli were presented on a grey background (RGB=128, 128, 128; see *Deviations from pre-registered methods*, 2). A fixation target was presented at the centre of the screen throughout the trial. It consisted of a filled black circle (0.5° diameter) with a white cross inside (width: 0.15°). To signal trial onset and to encourage fixation, a black dot (0.15° diameter) appeared inside the fixation target 300 ms before the onset of the memory array. The memory array consisted of two oriented, coloured bars inside circles (diameter: 6°, width: 0.3°). The colours were picked at random without replacement out of four possible colours on each trial: red (RGB=153, 0, 27), purple (RGB=122, 39, 150), green (RGB=16, 83, 0), and blue (RGB=0, 72, 136), chosen to be isoluminant in CieLab space. On every trial, the orientation of each stimulus was randomly selected without replacement from a uniform distribution of 160 orientation angles between 1.1250 and 180 degrees. The two WM items were presented laterally 6° from the fixation target. For the incidental task, participants were presented with a coloured ring around the fixation target (diameter: 1°, width: 0.5°). After the memory delay, the probe was a tilted bar inside a circle (6°; same size as memory items) and matched the colour of the probed memory item. The orientation angle of the probe was selected at random on each trial between 1 and 180 degrees. The probe was presented 6° above or below the fixation target, with equal probability, pseudorandomised for the full experimental session. This type of probe display has been reported to reduce the risk of stimulus-specific eye-movements (Mostert et al., 2018). Feedback of the correct target orientation was presented as a white line on top of their reported orientation for comparison.

### EEG data acquisition

Electroencephalography (EEG) was recorded at a sampling rate of 1kHz using Synamps amplifiers (Neuroscan, Inc., Albany, CA) and Curry Neuroimaging Suite (v7.0.12 X) software. Sixty-one electrodes were positioned according to the 10-10 system (American Clinical Neurophysiology Society, 1991). Eye movements were recorded from bipolar EOG derivations from electrodes above and below the right eye (vertical) and lateral to each eye (horizontal). A ground electrode was placed on the left elbow and the left mastoid was used as the active reference.

In addition, participants’ eye movements were recorded using an eye tracking camera (EyeLink) tracking at least one eye at a rate of 1kHz. Eye-movements from six participants were accidentally recorded at a sampling rate of 500 Hz instead of 1000 Hz. These participants were not excluded as the eye-tracking data was down-sampled to 500 Hz for all participants, in line with the pre-registered pre-processing pipeline.

### Pre-processing of eye-movement data

The EyeLink EDF file was converted to ASCII format and read into Matlab using the Fieldtrip toolbox (Oostenveld, Fries, Maris, & Schoffelen, 2011). The data were epoched around the onset of the WM array (−1 to 4 seconds). We took the mean of the right and the left eye (or used a single eye, where it was only possible to record from one eye; see *Deviations from pre-registered methods*, 3) and down-sampled the data to 500 Hz. Eye-blinks were detected (zeros in the eye-position data) and interpolated from -100 to +100 ms around the blink onset and offset using linear interpolation. Saccades (including microsaccades) were detected using a Matlab function (Engbert & Mergenthaler, 2006) based on two-dimensional velocity vectors computed from eye-position data using moving averages (velocity type=2). We used a velocity threshold (*λ*) of 6 standard deviations of the velocity distribution, and minimum duration of 6 ms.

### EEG Pre-processing

Pre-processing of the EEG data was performed in Matlab using the Fieldtrip toolbox (Oostenveld et al., 2011). The ERPs were re-referenced offline to the mean of the right and left mastoid and down-sampled to 500 Hz. A high-pass filter of 0.1 Hz and a low-pass filter of 40 Hz was applied to all channels. The data were epoched around the onset of the WM array (−1 to 4 seconds). We performed ICA to remove any heartbeat artefacts and/or artefacts due to eye-blinks. Artefact detection and rejection was performed separately for the time-window following WM array onset (0-1 s) and following cue onset (0-1.5 s). In addition, for an exploratory analysis not included in the pre-registration, we performed artefact rejection separately for a time-window following probe onset (0-1.5s). When an artefact was detected, we rejected the epoch containing the artefact from analyses, not the entire trial. We marked epochs with a high degree of variance using the Fieldtrip summary plot for visual artefact rejection and further marked epochs that exceed a cut-off threshold of 50 of the z-transformed value of the pre-processed data to detect any large artefacts due to muscle movements. We further excluded epochs with saccades >1° visual angle identified from the eye-tracking data (except for the probe epoch, as the probe was presented off-centre). In case eye-tracking data was missing, we visually inspected EOGs to reject saccade epochs. Six participants had missing eye-tracking data in the range of 4-32 trials.

We excluded participants with fewer than 80% of epochs remaining following artefact rejection from further analyses. On average, there were 1056.8 trials remaining for the encoding epoch analysis (min=944, max=1113), 1022.7 trials remaining for the cue epoch (min=863, max=1112) and 1097.3 trials remaining for the probe epoch (congruent: M=548.42, min=519, max=560; incongruent: M=548.88, min=517, max=559).

## Experimental design and statistical analyses

### Behaviour

The behavioural measures of interest were mean absolute error from the target angle and median response initiation times (henceforth RT). If selecting the cued colour during the maintenance delay privileges associated but currently irrelevant item-features, subsequent orientation judgements should be faster and more precise on congruent than on incongruent trials. This would replicate previous behavioural results (Zokaei, Manohar, et al., 2014; Zokaei, Ning, et al., 2014). Alternatively, if it is possible to selectively attend the cued colour in WM without strengthening memory representations for associated features of the same object, there should be no difference in performance between congruent and incongruent trials. We performed paired samples t-tests (one-tailed) to test for a main effect of congruency for both measures. We predicted that mean absolute error and median RTs would be significantly lower for congruent relative to incongruent trials.

To investigate sources of error in memory judgements and how these were affected by cueing, we fit a swap model to the data (Bays, Catalao, & Husain, 2009). The model is an extension of a mixture model (Zhang & Luck, 2008), which aims to identify the frequency of recalling the probed item, erroneously recalling the other item (‘swaps’), and random guessing. It can be described in the following way:

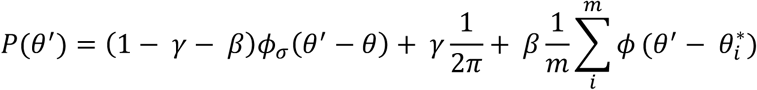

Where *θ* denotes the target orientation in radians, *θ*^′^ denotes the reported orientation, *γ* is the proportion of trials where the participant responds at random and *ϕ*_*σ*_ denotes the circular equivalent of the normal distribution (Von Mises distribution) with mean = 0 and standard deviation = *σ*. The swap model extension also includes the *β* parameter, which is the probability of responding with one of the non-target values: 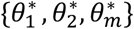 for m non-targets. As such, the model estimates the precision, target rate, non-target rate and guess rate of responses. As above, we performed paired samples t-tests to test for a main effect of congruency for each measure (one-tailed; congruent>incongruent for target rate and precision, and incongruent>congruent for guess rate and non-target rate; see *Deviations from pre-registered methods*, 4).

### Orientation decoding from EEG voltages

If selection of the cued colour reactivates other features of the cued item even though these are irrelevant to the incidental task, we expected to be able to decode the orientation of the cued item from EEG activity patterns following cue onset and decoding quality to be greater for the cued than the uncued item’s orientation. Alternatively, it may be possible to selectively access a single feature in memory, similar to selective coding for only relevant item features observed during encoding (Bocincova & Johnson, 2019; Serences et al., 2009). If so, patterns of activity in the EEG channels should contain no information about the cued item’s orientation and/or contain equal information about the orientations of both items following the cue.

To test whether the patterns of activity in the EEG channels contained information about the orientation of the items in memory, we used Mahalanobis distance to calculate the trial-wise distances between the multidimensional activity at each time point for each of the 160 possible orientations in memory, grouped into 16 angle bins distributed around a 180 degree angle space. The left and right memory items were decoded separately and independently for each participant. We performed the analyses on the 17 posterior channels (P7, P5, P3, P1, Pz, P2, P4, P6, P8, PO7, PO3, POz, PO4, PO8, O1, Oz and O2) as in (Wolff et al., 2017). Pre-stimulus baseline correction was performed by subtracting the mean voltage prior to WM array onset (−0.2 to -0.1 s) from voltages at all other time points.

The decoding procedure followed an 8-fold cross-validation approach to calculate the decoding accuracy of the orientation of interest for each trial. The activity pattern of the trials of the testing fold at a particular time-point were compared to the trials of the 7 training folds. The trials of the training folds were averaged into 16 orientation bins relative to the test trial orientation, each bin containing trials with orientations within a range of 11.25°. As trials were rejected due to artefacts, we ensured equal trial numbers in each bin by random subsampling. We computed the pairwise Mahalanobis distances between the test trials and each of the orientation bins using the covariance matrix estimated from trials in the 7 training folds, using a shrinkage estimator (Ledoit & Wolf, 2004). To obtain a visual representation of a tuning curve, the 16 distances were ordered as a function of orientation difference, mean-centered, and sign-reversed, so higher values represented greater similarity between the test trial and the training set. This was repeated for all train and test fold combinations. To obtain reliable estimates, the above procedure was repeated 100 times (with random folds each time), separately for eight orientation spaces used for binning the training trials with respect to the test trial (bin centres: 1.41°-170.16°, 2.81°-171.56°, 4.22°-172.97°, 5.63°-174.38°, 7.03°-175.78°, 8.44°-177.19°, 9.84°-178.59° and 11.25°-180°, each in steps of 11.25°). The resulting 800 samples (100 repetitions×8 orientation spaces) were averaged for each of the 16 Mahalanobis distance values at each time point.

Finally, to obtain a summary measure of decoding accuracy, we computed the cosine-weighted means of the tuning curves (Sprague, Ester, & Serences, 2016; Wolff et al., 2017). Higher values reflect greater orientation tuning and thereby greater decoding quality for that trial and chance-level is zero. We took the mean over trials, resulting in a single decoding value for each time-point for each participant.

To test for significant orientation decoding of the two items, we took the mean decoding accuracy within the time-window following array onset (0–1 s; WM array epoch) and following cue onset (0–1 s; cue epoch). As a positive control we tested if the orientation of both items could be decoded during the WM array epoch using one-sample t-tests (one-tailed>0) and tested for any differences in decoding strength between items using a paired sample t-test (two-tailed). We expected to be able to decode the orientation of both items during the WM array epoch and decoding quality to be similar for both items.

The primary analysis of interest tested whether the orientation of the cued item could be decoded during the cue epoch. Again, we used one sample t-tests, to test for significant decoding of each item (one-tailed, >0) and a paired sample t-test (one-tailed, cued>uncued) to test for any difference in decoding for the two items. We expected orientation decoding to be significant for the cued item and decoding quality to be significantly greater for the cued than uncued item.

For completeness, we also tested for significant decoding of each memory item and for any difference between them at each time-point (one-sample t-test, one-tailed, >0, paired samples t-test, two-tailed) using a cluster-based permutation test (10,000 permutations) to correct for multiple comparisons over time. We used a cluster-forming and cluster significance threshold of *p*<.05. Time-resolved decoding values were smoothed with a Gaussian kernel (SD=20 ms) for visualisation and statistical tests.

### Spatiotemporal decoding of orientation from EEG

Decoding accuracy may be improved by taking advantage of the dynamic nature of ERPs (Grootswagers, Wardle, & Carlson, 2016; Wolff, Jochim, Akyürek, Buschman, & Stokes, 2020). This approach pools the relative voltages over time and sensors, thus leveraging information encoded in temporal as well as spatial patterns. Building on the pre-registered methods, we complement the standard decoding with this spatiotemporal decoding approach focusing on a 300 ms time window from 100-400 ms following onset of the WM array and following the cue, consistent with a recent similar study (Wolff et al., 2020). Pre-stimulus baseline correction (- to -0.1 s before WM array onset) was performed on the raw voltages as described above. The data were then down-sampled to 100 Hz by taking the average every 10 ms. The resulting 30 values for each posterior channel were concatenated and used as the input to the multivariate decoder. The decoding procedure followed the same steps described above, but resulted in a single decoding value for each time-window of interest. As above, the procedure was repeated 100 times (with random folds each time), separately for eight orientation spaces and averaged across the resulting 800 samples for each of the 16 Mahalanobis distance values at each time-point.

As above, we used one sample t-tests, to test for significant decoding of each item (one-tailed, >0). During the WM array epoch, we expected decoding strength to be significant for both items. During the cue epoch, we expected orientation decoding to be significant for the cued item and decoding strength to be greater for the cued than the uncued item (one-tailed; cued>uncued).

### Time frequency decomposition and alpha lateralization

Using alpha lateralization and gaze bias as indices of spatial attention in WM, we tested if the centrally presented, non-predictive colour cue engaged spatial attention toward the original position of the cued item, despite location being a fully redundant feature in this task.

If so, we expected contralateral minus ipsilateral alpha power to be reduced relative to the original location of the cued item. We computed the alpha-band power lateralization index relative to the cued item’s location in the time window after incidental cue onset (0.2–0.8 s; to avoid effects related to probe onset). Spectral decomposition of the data was performed in Matlab with the FieldTrip toolbox (Oostenveld et al., 2011). We applied Hanning tapers with a time window width of five cycles per frequency, with frequencies of interest between 8 Hz and 14 Hz in steps of 1 Hz. We log-transformed the power at each frequency 10 × *log*_10_ and calculated the mean power over contralateral and ipsilateral posterior channels (P8/7, P6/5, P4/3, PO8/7, PO4/3, O2/1) relative to the cued item. To get a lateralization index, we took the difference between contralateral and ipsilateral alpha power averaged over the specified time-window (0.2-0.8s) and tested for significant lateralization using a one-sample t-test (one-tailed, <0; see *Deviations from pre-registered methods*, 5). In an exploratory analysis, we also tested for significant alpha lateralization at each time-point across the time-window from 0-1 s (one-sample t-test, two-tailed) using a cluster-based permutation test (10,000 permutations) to correct for multiple comparisons over time. We used a cluster-forming and cluster significance threshold of *p*<.05.

### Gaze bias analysis

Human gaze may be biased toward attended spatial locations in WM (Van Ede, Board, & Nobre, 2020; van Ede et al., 2019). Here, we similarly used gaze bias as an index of spatial attention towards the original position of the cued item. If spatial attention is directed toward the original position of the cued item, we expected horizontal gaze position to be biased toward the left (i.e., negative values) for left cue trials and toward the right (i.e., positive values) for right cue trials. First, we centred the gaze position data to the mean position during fixation (300–100 ms before stimulus onset). We then normalized the data according to the eccentricity of the memory items, so +/− 100% corresponds to the eyes being focused on the original position of the memory items (i.e., 6° visual angle) and 0% corresponds to fixation. We plotted the gaze position following incidental cue onset separately for left and right cue trials. Heat maps of gaze density were created by computing the 2D histograms (bin size=.012 dva) of eye-position from 0.2-0.8 s from cue onset separately for the two possible locations of the cued memory item. We converted histogram counts to density by dividing by the trial number for each location condition before subtracting out density common to both locations. Finally, we took the mean across participants (see *Deviations from pre-registered methods*, 6).

To obtain an aggregate measure of ‘towardness’, we took the mean horizontal gaze position for right cue trials minus the mean gaze position for left cue trials, divided by two, averaged across the time-window following incidental cue onset (0.2–0.8) (van Ede et al., 2019). We tested for significant gaze bias toward the cued item using a one-sample t-test (one-tailed, >0). In an exploratory analysis, we also tested for significant towardness at each time-point across the time-window from 0-1 s (one-sample t-test, two-tailed) using a cluster-based permutation test (10,000 permutations) to correct for multiple comparisons over time. We used a cluster-forming and cluster significance threshold of *p*<.05. Time-resolved towardness values were smoothed with a Gaussian kernel (SD=20 ms) for visualisation and statistical tests.

### Deviations from pre-registered methods

1. We had planned to recruit a total of 35 participants, but due to Covid-19 restrictions only 32 participants completed the experiment and 26 participants were included in the analyses following exclusion as per the pre-registered criteria.
2. The pre-registration mistakenly stated stimuli would be presented on a grey background with RGB values: 0.5, 0.5, 0.5, when the correct RGB values are: 128, 128, 128.
3. The pre-registration stated we would take the mean of the right and the left eye position. Due to some participants wearing glasses, it was only possible to record from one eye. We did not exclude these participants, but used the position data from the single recorded eye in these cases.
4. When testing for an effect of congruency on the parameters of the mixture model, the pre-registration mistakenly stated ‘two-tailed’ paired samples t-tests would be performed. As we specifically predicted performance to be better for congruent than incongruent trials, we performed one-tailed paired samples t-tests to test for a main effect of congruency for each measure (congruent>incongruent for target rate and precision, and incongruent>congruent for guess rate and non-target rate).
5. When testing for significant alpha lateralization following cue onset, the pre-registration specified the wrong direction of the one-tailed test (‘>0’ should be ‘<0’). We were specifically interested in contralateral alpha suppression, so we expected contra– ipsilateral alpha power to be negative.
6. The precise methods for plotting the eye-position heat maps were not included in the pre-registration, but are included in the methods for clarity.

### Exploratory analyses

#### Relationship between cue-locked orientation decoding and behaviour (error/RT)

Representational quality of the cued item’s orientation may signal the extent to which the cued item enters a privileged state in WM. If so, greater orientation decoding quality may be associated with better performance on congruent trials and worse performance on incongruent trials. We tested for a trial-wise relationship between orientation decoding quality and behavioural performance. Two general linear models were fitted for each participant with trial-wise orientation decoding quality averaged over the time-window of interest (0-1 s from cue onset) as the predictor variable. The outcome variable was log-transformed RTs in one model and absolute error in the other. The design matrices included a constant term. We fit the models separately for congruent and incongruent trials and obtained *β* weights for each participant. We performed one-sample t-tests on the *β* values. We expected greater decoding quality following the cue to predict lower RT/error on congruent trials (one-tailed, <0) and higher RT on incongruent trials (one-tailed, <0).

To test whether potential relationships to behaviour were specific to particular time-points in the entire post-cue time-window, we fit the regression models over time to obtain *β* weights for each time point during the delay following cue onset and performed cluster-corrected t-tests across a time-window from 0-1.5 s following cue onset (two-tailed, cluster-forming threshold=.05, 10,000 permutations). Time-resolved *β* values were smoothed with a Gaussian kernel (SD=20 ms) for visualisation and statistical tests.

#### Relationship between cue-locked alpha lateralization/gaze bias and behaviour (error/RT)

We tested for a trial-wise relationship between alpha lateralization/gaze bias and behavioural orientation recall performance (error/RT) using the same methods as described for orientation decoding above, but for the time-window of interest from 0.2-0.8 s following cue onset. We performed one-sample t-tests on the trial-wise *β* values. We expected contralateral alpha suppression to predict lower RT/error on congruent trials (one-tailed, >0) and higher RT/error on incongruent trials (one-tailed, >0). We expected greater gaze bias following the cue to predict lower RT/error on congruent trials (one-tailed, <0) and higher RT on incongruent trials (one-tailed, <0). As for orientation decoding, to test if any relationships to behaviour were specific to particular time-points in the post-cue delay, we also fit the regression models over time and performed cluster-corrected t-tests across the entire post-cue period (0-1.5 s following cue onset).

#### Probe processing

At orientation recall, participants compared the orientation of the randomly oriented probe stimulus to the probed item in memory in order to turn the dial to the appropriate orientation angle (target). The neural representation of the absolute distance between the random probe orientation and the memory item may signal how well the memory item is compared to the probe at recall. In this analysis, we aimed to test whether computation of this comparison signal was modulated by whether the target item for recall was previously cued (congruent) or not (incongruent).

#### Decoding target-probe distance prior to response onset

We performed the Mahalanobis distance decoding procedure specified above on the absolute angular distance between the target item in memory and the randomly oriented probe presented at orientation recall (target-probe distance). Instead of sorting the trials into 16 bins from 0-180°, the trials were averaged into 8 angular distance bins from 0-90°, each bin containing trials with orientations within a range of 11.25°. Otherwise the decoding procedure followed the same 8-fold cross-validation approach to calculate the decoding accuracy of the absolute angular distance of interest for each trial, repeated 100 times with random folds each time. Finally, as the absolute angular distance had a bounded uniform distribution rather than circular distribution, we computed the linear slope of the tuning curve instead of the cosine-weighted means to obtain a summary measure of decoding quality (Muhle-Karbe et al., 2021).

Training and testing was done separately for congruent and incongruent trials. The analysis was performed on the time-window from –0.5-0 s before the response onset (response-locked). Since reaction times were mostly longer than 0.5 s (72.2% of trials), the response-locked epoch mostly contained time points after the onset of the probe (Figure 5G). One-sample t-tests were performed to test for significant target-probe distance decoding for congruent and incongruent trials within in time-window of interest (–0.5-0s; one-tailed, >0) and a paired-samples t-test was performed to test for a significant difference between congruent and incongruent trials (two-tailed). Additionally, cluster-corrected t-tests were performed across the full time-window prior to response onset (–1-0 s; two-tailed, cluster-forming threshold=.05, 10,000 permutations). Decoding time courses were smoothed with a Gaussian kernel (SD=20 ms) for visualisation and statistical tests.

#### Decoding target-probe distance and nontarget-probe distance following probe onset

To preview the results, we found significant decoding of the target-probe distance prior to response onset and decoding quality was greater on congruent than incongruent trials. We next asked whether, on incongruent trials, the probe might initially be compared to the cued item, even though it is the non-target. To capture the probe-evoked neural response, the analysis was time-locked to the probe-onset (0-0.5s) rather than to response onset as above. Using the same decoding procedure described above, we decoded the target-probe distance and the nontarget-probe distance. Note that on congruent trials, the cued item is the same as the target, whereas on incongruent trials, the cued item is the nontarget. Thus, decoding strength of the nontarget-probe distance on incongruent trials may indicate the extent to which the cued item is incorrectly used as a template for comparison with the probe.

One-sample t-tests were performed to test for significant decoding of the target-probe and nontarget-probe distance for congruent and incongruent trials and for significant differences between them within the time-window of interest (0-0.5s; one-tailed, >0). Cluster-corrected t-tests were performed across the full time-window following probe onset (0-0.5s; two-tailed, cluster-forming threshold=.05, 10,000 permutations). Time-resolved decoding values were smoothed with a Gaussian kernel (SD=20 ms) for visualisation and statistical tests.

Significant nontarget-probe distance coding could be a generic comparison that is unrelated to performing the task and preparing a response. To test the extent to which the neural pattern coding for the nontarget-probe distance resembled the neural pattern coding for the target-probe distance on congruent trials, we repeated the same analysis, but instead of training and testing the decoder within each condition, the training data were congruent trials labelled with target-probe distance values, while the test data were either congruent or incongruent trials labelled with non-target probe distance values. By training on the same data (target-probe distance), we have a better chance of comparing the same neural mechanism. On congruent test trials, 8-fold cross validation was performed as above. As the training and test data were independent on incongruent test trials, we did not perform 8-fold cross-validation. Significant decoding in this analysis would indicate the neural pattern coding for the nontarget-probe distance resembles the neural pattern coding for the target-probe distance when the target was previously cued.

#### Relationship between probe processing and behavioural error

To test if target-probe / nontarget-probe distance decoding quality was related to behavioural performance on the orientation recall task, we fit a general linear model with EEG decoding values averaged over the time-window of interest (response-locked: –0.5-0s; probe-locked: 0-0.5s) as the predictor variable and absolute error as the outcome variable. The design matrix included a constant term. We fit the models separately for congruent and incongruent trials. We performed one-sample t-tests on the *β* weights to test whether the quality of decision-signal decoding was associated absolute error. We expected target-probe distance decoding quality to be associated with reduced error (one-tailed, <0) and nontarget-probe distance decoding quality to be associated with greater error (one-tailed, >0).

## Results

### Orientation recall is faster and more precise on congruent relative to incongruent trials

Participants performed well on the incidental task with mean accuracy of .972 (SD=.021) and mean median RT on correct trials was .592 s (SD=.052). Correct incidental task trials were included in the subsequent analyses.

Behavioural orientation recall data are shown in Figure 1B-C. As expected, mean absolute error was significantly lower for congruent (M=13.75, SD=3.22) compared to incongruent trials (M=20.67, SD=6.08): *t*_25_=-7.01, *p*<.001, *d*=-1.38 (Figure 1C, left panel). Similarly, RTs were significantly lower for congruent (M=.55, SD=.12) compared to incongruent trials (M=.67, SD=.15): *t*_25_=-8.35, *p*<.001, *d*=-1.64 (Figure 1C, right panel). Selecting the cued colour in the incidental task facilitated speed and accuracy of subsequent recall of other features belonging to the same item relative to the other item in memory.

### Congruency affects all sources of error

Errors in orientation judgments may arise from multiple sources including reduced precision, reduced probability of reporting the correct memory item (target rate), increased likelihood of guessing (guess rate) and/or increased likelihood of reporting the other memory item (non-target rate). The estimated parameters of the swap model are shown in Figure 1D. Congruency affected all sources of error. Precision (K) was significantly higher for congruent (M=4.50, SD=1.30) than incongruent (M=3.27, SD=1.27) trials: *t*_25_=6.31, *p*<.001, *d*=1.24. The same was true for the target rate (congruent: M=.94, SD=.06; incongruent: M=.79, SD=.14): *t*_25_=5.74, *p*<.001, *d*=1.13. Guess rates were significantly lower for congruent (M=.05, SD=.05) than incongruent (M=.17, SD=.13) trials: *t*_25_=-4.73, *p*<.001, *d*=-.927, and so was the non-target rate (congruent: M=.02, SD =.02; incongruent: M=.04, SD=.06): *t*_25_=-2.20, *p*=.019, *d*=-.431.

### Non-predictive colour cue strengthens the neural representation of the cued item’s orientation

If attentional selection of the cued colour for the incidental task spreads to other features belonging to the cued item, we would expect the neural representation of the cued item’s orientation to be reinstated following the cue.

First, we ensured that the orientation of both items could be decoded during the WM array epoch (Figure 2A). Orientation decoding quality was significant for both memory items (cued: *t*_25_=4.55, *p*<.001, *d*=.894; 0.07-0.76 s from WM array onset, cluster-corrected *p*<.001; uncued: *t*_25_=5.31, *p*<.001, *d*=1.04; 0.06-0.80 s from WM array onset, cluster-corrected *p*<.001) and there was no significant difference between them (*p*=.780). We use labels ‘cued’ and ‘uncued’ to keep consistent across analyses even though the cue does not appear until later in the trial. Note that decoding quality dropped to near baseline prior to cue onset.

**Figure 2.**
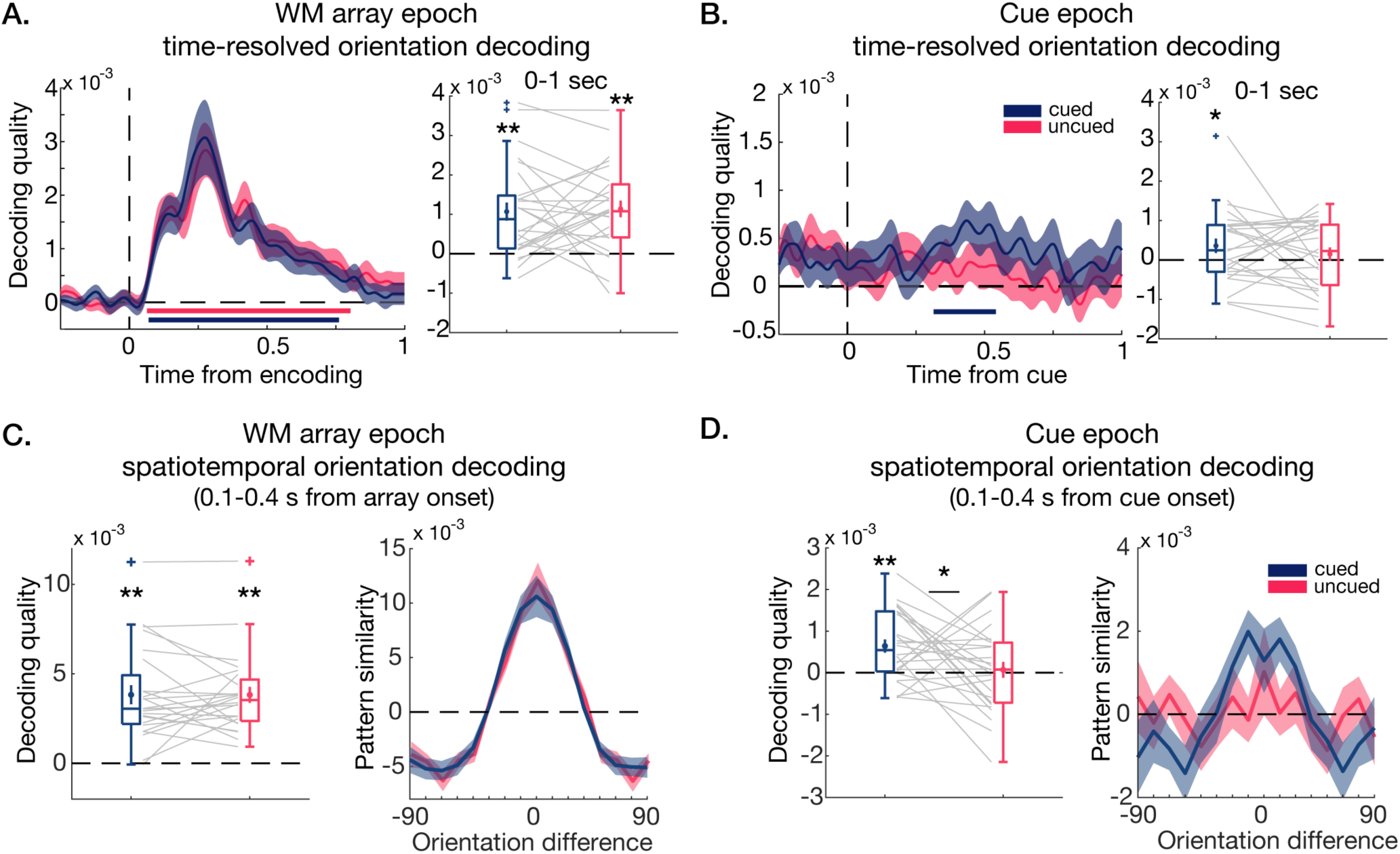
Decoding orientation from EEG voltages. **A**. Left: Decoding quality following WM array onset. Shaded area shows SEM. Bars show clusters of significant decoding. Boxplot shows mean decoding across the encoding epoch (0-1s from WM array onset). Overlaid mean and SEM. Grey lines show individual participants. **B**. Same as A but for cue epoch. **C**. Spatiotemporal orientation decoding. Left: Boxplot of decoding quality following WM array onset (100-400 ms). Right: Mahalanobis distance pattern similarity as a function of orientation difference. Shaded area shows SEM. **D**. Same as C but following cue onset (100-400 ms).

Our main analysis of interest tested whether decoding quality for the cued item’s orientation re-emerged following cue onset (Figure 2B). As predicted, decoding quality for the cued item’s orientation was significantly greater than zero in the cue epoch (*t*_25_=1.93, *p*=.032, *d*=.379; 0.30-0.54 s after the cue, cluster-corrected *p*=.025), while the orientation of the uncued item could not be decoded (*p*=.20). However, there was no significant difference in decoding quality between the cued and uncued item (*p*=.18). The results suggest that attentional selection of the cued colour strengthens the neural representation for the cued item’s orientation, even though the item’s orientation is irrelevant at the time of the cue.

In an exploratory analysis, we applied a complementary spatiotemporal decoding approach to boost power to detect neural patterns coding for memorised orientations using EEG (Figure 2C-D). Confirming the standard decoding results, the orientations of both items could be decoded following WM array onset (100-400 ms; cued: *t*_25_=7.47, *p*<.001, *d*=1.46; uncued: *t*_25_=8.78, *p*<.001, *d*=1.72), and there was no difference in decoding strength between the two items (*p*=.94).

In the cue-locked analysis of interest (100-400 ms after cue onset) the orientation of the cued item, but not the uncued item, could be decoded (cued: *t*_25_=3.96, *p*<.001, *d*=.776; uncued: *p*=.361) and decoding strength was significantly higher for the cued item’s orientation compared to the uncued item (*t*_25_=2.17, *p*=.020, *d*=.425). Thus, the results of the spatiotemporal decoding analysis broadly confirm the standard decoding results and further suggest that the cue-induced representational enhancement is greater for the cued than the uncued item, even though the cued item’s orientation was no more relevant to the task than the uncued item’s orientation.

The EEG-based orientation decoding results were not explained by fixational eye-movements (Mostert et al., 2018; Quax, Dijkstra, van Staveren, Bosch, & van Gerven, 2019). We repeated the same orientation decoding analysis on eye-position data instead of EEG data (Figure 3A-B). Decoding from eye-position was not quite significant for the cued item’s orientation following cue onset, but was significant for the uncued item’s position (cued: *t*_25_=1.63, *p*=.057, *d*=.321; uncued: *t*_25_=1.81, *p*=.041, *d*=.355; difference: *p*=.35). However, two participants were outliers and showed strong eye-based orientation decoding for the cued item. Thus, we repeated the EEG decoding analyses excluding these participants to show that EEG-based orientation decoding was not driven by participants with strong decoding from eye-position (Figure 3C-D).

**Figure 3.**
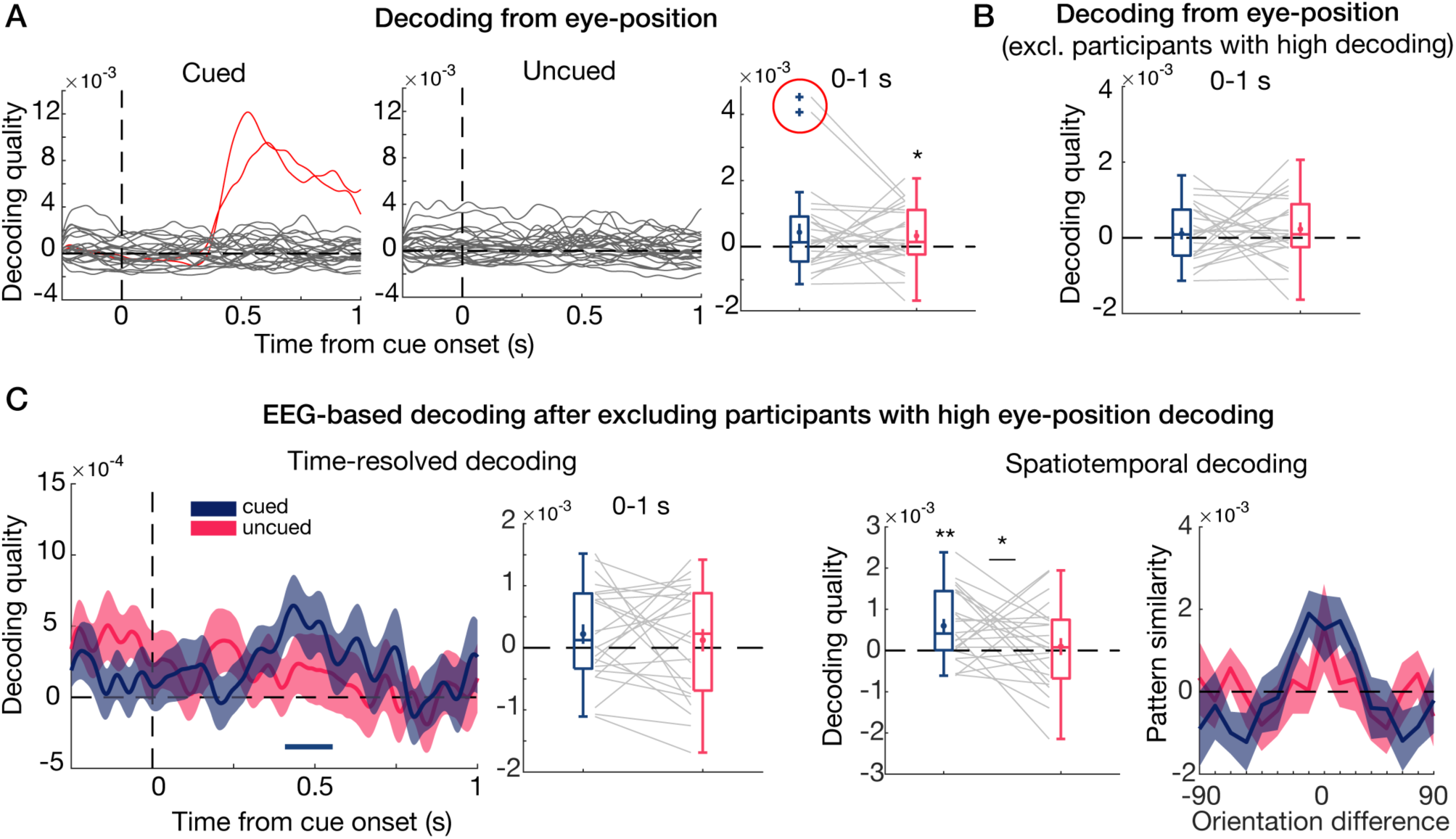
Controlling for fixational eye-movements. **A**. Eye-position based decoding of orientation. Decoding quality over time for all participants (individual lines) for cued (left) and uncued (right) item. Boxplot shows mean decoding quality across 0-1s from cue onset. Overlaid mean and SEM. Grey lines show individual participants. Red lines and circle represent two outliers excluded from control analyses. **B**. Boxplot showing mean eye-position based orientation decoding after excluding participants with high eye-position decoding. **C**. EEG-based decoding of orientation after excluding participants with high eye-position decoding. Left: standard time-resolved decoding quality. Blue bar represents significant time-points after cluster-correction (0.38-0.53 s., *p*=.053). Mean decoding (cued: *t*_23_=1.40, *p*=.088, *d*=.285; uncued: *p*=.25; difference: *p*=.32). Right: spatiotemporal decoding quality (cued: *t*_23_=3.51, *p*<.001, *d*=.717; uncued: *t*_23_=.46, *p*=.32, *d=*.095; difference: *t*_23_=1.83, *p*=.040, *d*=.373). \**p*<.05, \*\**p<*.001.

### No evidence for relationship between orientation decoding quality and behaviour

If the observed representational enhancement of the cued item’s orientation strengthens the memory quality and accessibility of the cued item relative to the uncued item, trial-wise variance in decoding quality for the cued item’s orientation may predict speed and accuracy of subsequent orientation recall. However, we found no evidence for a trial-wise relationship between post-cue orientation decoding in the time-window of interest (0-1 s) and behavioural measures on either congruent (RT: *p*=.49, absolute error: *p*=.81) or incongruent trials (RT: *p*=.84; absolute error: *p*=.61). There were also no significant clusters in the time-resolved regression across the entire post-cue delay (0-1.5 s). Similarly, there was no relationship between spatiotemporal decoding quality and behavioural measures on congruent (RT: *p*=.64, absolute error: *p*=.75) or incongruent trials (RT: *p*=.50, absolute error: *p*=.43).

### Colour cue triggers alpha suppression contralateral to cued item’s memorised location

Next, we asked if selecting the cued colour for the incidental task engaged spatial attention toward the original position of the cued item, even though location information was irrelevant to the task. If so, we expected to see alpha suppression contralateral to the cued item’s original position. Figure 4A shows contralateral minus ipsilateral power across a range of frequencies following cue onset, and Figure 4B shows mean contra-minus ipsilateral alpha power (8-14Hz). As predicted, contralateral alpha power was significantly lower than ipsilateral alpha power in the time-window following cue onset (0.2-0.8 s): *t*_25_=-4.78, *p*<.001, *d*=-.938. Time-resolved alpha lateralization following cue onset (0-1 s) was significant from 0.22 to 0.71 s (cluster-corrected *p*<.001). Thus, colour-selection in WM is associated with transient suppression in contralateral alpha power, even when memory for location is not demanded by the task.

**Figure 4.**
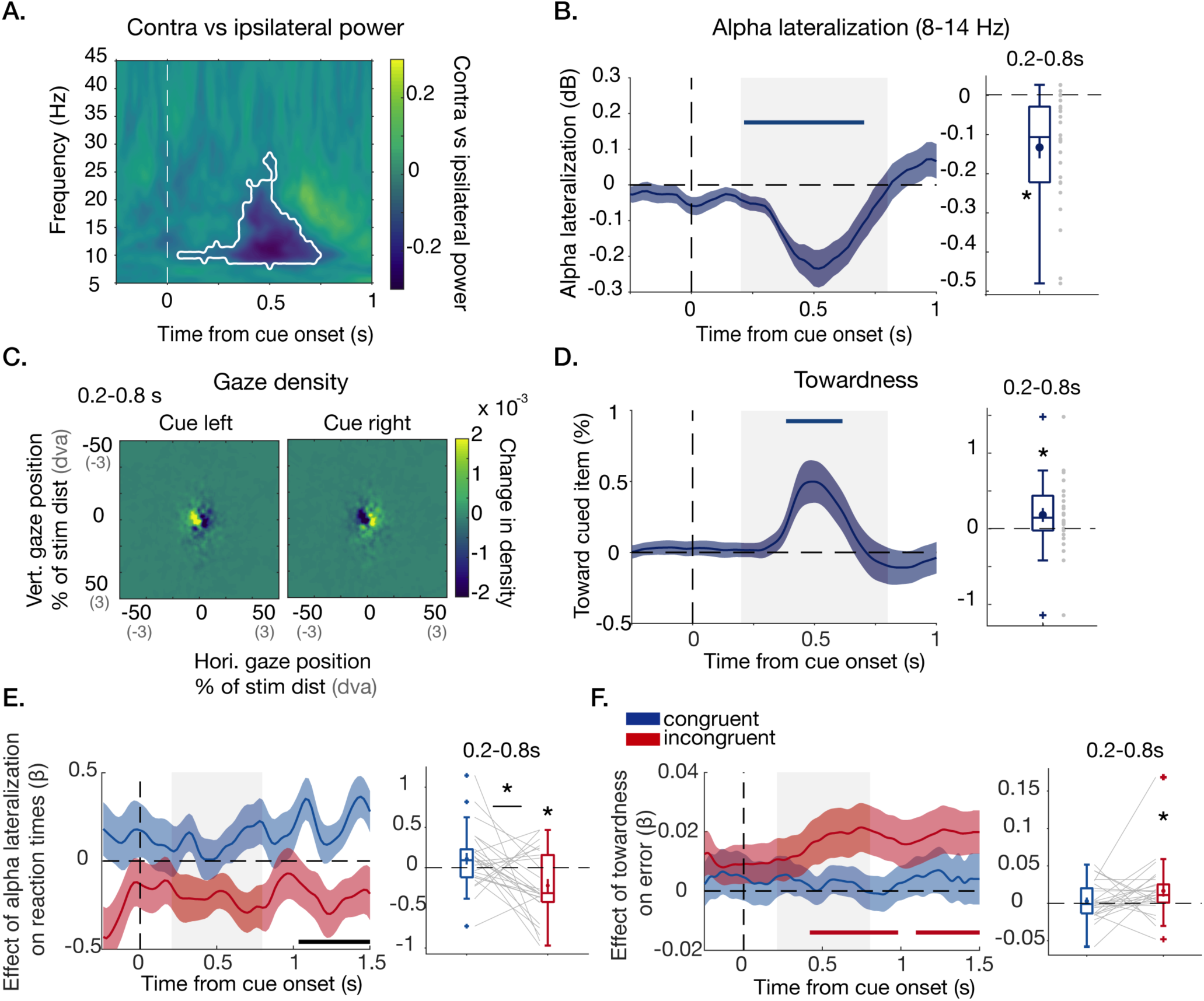
Alpha lateralization and gaze bias **A**. Contra minus ipsilateral power relative to original position of cued item over time from cue onset. White contours show cluster of significant contralateral suppression. **B**. Left: alpha lateralization (contra minus ipsilateral alpha power; 8-14 Hz) over time from cue onset. Grey shaded area indicates time window of interest (0.2-0.9 s). Shaded area shows SEM. Right: Boxplot of mean alpha lateralization across cue epoch (0.2-0.8 s from cue onset). Mean and SEM overlaid. Grey dots show individual participants. **C**. Two-dimensional histogram of gaze density for cue left and cue right trials (subtracting mean gaze across conditions). **D**. Mean horizontal gaze position for right cue trials minus the mean gaze position for left cue trials, divided by two (“towardness”) over time from cue onset. Positive values indicate gaze bias toward cued item. Right: Boxplot of mean towardness across cue epoch (0.2-0.8 s from cue onset). **E**. Relationship between alpha lateralization and reaction times. Regression weights (β) from trial-wise regression with alpha lateralization and reaction times for congruent (blue) and incongruent (red) trials. Left: β weights from time-resolved regression. Right: β weights from regression with mean alpha lateralization across cue epoch. Black bar shows significant difference between congruent and incongruent trials. **F**. Same as E for relationship between gaze bias and absolute error. \**p*<.05.

### Colour cue triggers gaze bias toward cued item’s memorised location

Eye-position data are shown in Figure 4C-D. As predicted, mean gaze position was significantly biased toward the original position of the cued item as indicated by towardness values greater than zero (M=.186%, SD=.495%) following cue onset (0.2-0.8s): *t*_25_=1.91, *p*=.034, *d*=.375 (Figure 3D). Time-resolved towardness following incidental cue onset (0-1 s) was significant from 0.38 to 0.61 s (cluster-corrected *p*=.044), suggesting colour-selection triggers a transient shift in gaze position toward the cued item’s memorised position. It is worth noting that, while statistically robust, this bias was tiny on average (.186% of 6° = 0.011° visual angle). As also evident from the 2D histogram (4C), it did not arise from overt saccades to the item’s location on a proportion of trials, but rather small but consistent shifts of gaze position.

### Contralateral alpha suppression predicts speed of orientation recall

Next, we tested whether there was a trial-wise relationship between measures of spatial attention (contralateral alpha suppression and gaze bias) and behavioural performance (error and RT) in a set of exploratory analyses. If spatial attention toward the cued item’s original position is involved in the selection and prioritisation of the cued item’s other features in WM, contralateral alpha suppression and gaze bias should be associated with faster and/or more precise performance on congruent trials. If selection of the cued item comes at the expense of the uncued item in memory, alpha suppression and gaze bias might be associated with slowed and/or less precise performance on incongruent trials.

Trial-wise contralateral alpha suppression was associated with slowed RTs on incongruent trials (Figure 4E; *t*_25_=-2.73, *p*=.006, *d*=-.535). While β weights on congruent trials tended towards the opposite direction, contralateral alpha suppression did not significantly predict faster RTs on those trials (*t*_25_=1.49, *p*=.074, *d*=.292). There was a significant difference in βs between congruent and incongruent trials (*t*_25_=2.72, *p*=.012, *d*=.534; 1.04 to 1.5 s cluster-corrected *p*=.005). There was no evidence of a trial-wise relationship between contralateral alpha suppression and absolute error (congruent: *p*=.456; incongruent: *p*=.357; difference: *p*=.809).

Greater suppression of alpha contralateral to the cued item’s original position predicted slowed response initiation when the other memory item is subsequently probed for orientation recall, suggesting contralateral alpha suppression may reduce the accessibility of the uncued relative to the cued item in memory.

### Gaze bias is associated with greater error on incongruent trials

The trial-wise regression analyses showed that gaze bias toward the cued item’s original position was associated with greater absolute error on incongruent trials (*t*_25_= 2.16, *p*=.020, *d*=.424; 0.42 to 0.98 s cluster-corrected *p*=.023; 1.10 to 1.5 s, cluster-corrected *p*=.029). There was no significant relationship with gaze bias and absolute error on congruent trials (*p*=.72) and while β weights were larger for incongruent than congruent trials, this difference was not significant (*t*_25_=-1.53, *p*=.14, *d*=-.299). There was also no evidence of a trial-wise relationship between gaze bias and RT (congruent: *p*=.44; incongruent: *p*=.75; difference: *p*=.72). The results indicate that gaze bias toward the cued memory item’s memorized position may reduce memory quality for the other item.

In summary, both alpha lateralization and gaze bias significantly influenced behaviour, but in distinct ways. While alpha lateralization primarily predicted reaction times, gaze bias primarily predicted errors (specifically on incongruent trials).

### No correlation between alpha lateralization and gaze bias

There was no relationship between alpha lateralization and gaze bias following the cue (0.2-0.8s from cue onset) on a trial-wise (β weights not significantly different from zero, *p*=.60) or participant-wise level (*r*s_24_=-.264, *p*=.19).

### Target-probe comparison signal stronger on congruent trials

The results so far indicate that attentional selection of the cued colour during the delay is associated with faster and more precise recall of the cued relative to the uncued item’s orientation, enhances the neural representation of the cued item’s orientation and triggers spatial attention toward the cued item’s original position. Next, we investigated whether attentional selection of the cued item during the delay modulates how effectively the memory items are compared to the probe at recall in a set of exploratory analyses.

At recall, participants were required to compare the orientation of the randomly oriented probe stimulus to the probed item in memory (target) in order to turn the dial to the appropriate angle. The neural representation of the absolute distance between the random probe orientation and the target (target-probe distance) may signal how well the memory item is compared to the probe at recall. The absolute target-probe distance could be decoded prior to response onset (mean decoding -0.5 to 0 s) on both congruent (*t*_25_=6.16, *p*<.001, *d*=1.21; -0.68 to 0 s, cluster-correction *p* <.001) and incongruent trials (*t*_25_=2.95, *p*=.003, *d*=.579; -0.28 to 0 s, cluster-corrected *p*=.012; Figure 5A). Target-probe distance decoding quality was significantly higher on congruent than incongruent trials (*t*_25_=3.41, *p*=.002, *d*=.668; -0.40 to 0 s, cluster-corrected *p*=.002). Thus, the target-probe distance comparison signal was stronger prior to response onset when the item had previously been selected from memory.

**Figure 5.**
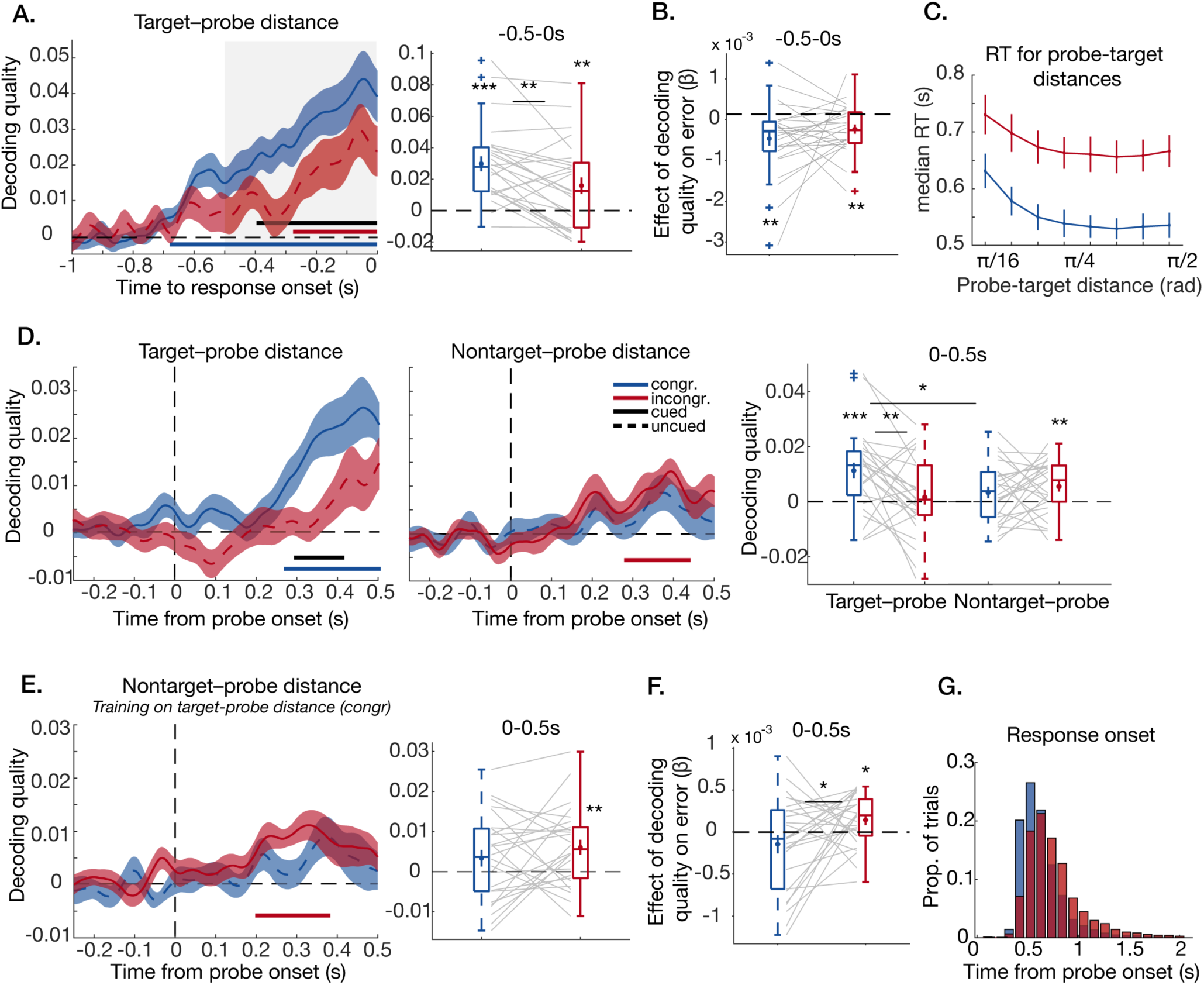
Decoding target-probe and nontarget-probe distance from EEG voltages. **A**. Decoding target-probe distance (absolute distance between random probe orientation and orientation of probed item in memory) for congruent (blue) and incongruent (red) trials time-locked to response onset. Shaded area shows SEM. Bars show significant decoding. Grey shaded area indicates time window of interest (−0.5– 0s). Boxplot (right) shows mean decoding strength before response onset (−0.5-0 s). Mean and SEM overlaid. Grey lines show individual participants. **B**. Relationship between target-probe distance decoding and orientation recall error. Regression weights (β) from trial-wise regression with mean probe-target distance decoding (−0.5-0s before response onset) and absolute error for congruent (blue) and incongruent (red) trials. **C**. Mean median reaction time (s) as a function of probe-target distance (in radians) on congruent (blue) and incongruent (red) trials. Error bars show SEM. **D**. Decoding of target-probe distance (left) and nontarget-probe distance (middle) for congruent (blue) and incongruent (red) trials time-locked to probe onset. Boxplot (right) shows mean decoding strength following probe onset (0-0.5s). Solid lines show the item that was cued during the delay and dashed lines show the uncued item. **E**. Same as middle panel in D, except decoding training data was sampled from congruent trials labelled with target-probe distance values. **F**. Relationship between nontarget-probe distance decoding (trained on congruent target-probe distance; shown in E) and orientation recall error. Regression weights (β) from trial-wise regression with mean nontarget-probe distance decoding (0-0.5 after probe onset) and absolute error. **G**. Histogram of response onset relative to probe onset across participants for congruent (blue) and incongruent (red) trials. \**p*<.05, \*\**p*<.01, \*\*\**p*<.001.

RTs varied as a function of probe-target distance, with slower RTs when the probe-target distance was small (Figure 5C). However, RT was not differently modulated by probe-target distances on congruent and incongruent trials (Figure 5C; 2-way ANOVA, interaction between congruency and probe-target distance, *F*_7,25_=1.31, *p*=.247), making it unlikely that the reported difference in probe-target distance decoding quality between congruent and incongruent trials is driven by differences in RT (or, equivalently, how recently the probe has appeared).

### Target-probe comparison signal predicts orientation recall error

We tested whether there was a trial-wise relationship between the target-probe distance signal prior to response onset and behavioural performance (Figure 5B). We found that target-probe distance decoding quality correlated with orientation recall error on both congruent (*t*_25_=-3.36, *p*=.001, *d*=-.660) and incongruent (*t*_25_=-3.17, *p*=.002, *d*=-.622) trials, indicating that a stronger representation of the probe-target distance magnitude predicted lower behavioural error in both cases. There was no difference between conditions (*p*=.29).

### Probe-locked decoding of target-probe and nontarget-probe distance

If the item selected for the incidental task during the delay is in a prioritised state for comparison with the probe at recall, we might expect a comparison signal between the probe and the *cued* item regardless of whether it is the target item probed for orientation recall, as is the case on congruent trials, or the nontarget item, as is the case on incongruent trials. This analysis focused on the probe-evoked response (0-0.5 s following probe onset). We would expect any erroneous comparison to the non-target to occur right after probe onset, especially since swap errors were very rare in both conditions (<5%), meaning that by the time participants start responding they have likely retrieved the target item.

The absolute distance between the probed memory item and the random probe orientation (target-probe distance) could be decoded following probe onset on congruent, but not on incongruent trials (congruent: *t*_25_=3.91, *p*<.001, *d*=.768; 0.26 to 0.5 s, cluster-corrected *p*<.001; incongruent: *p=*.27; Figure 5D). Mirroring the results of the response-locked analysis, target-probe distance decoding strength was significantly higher on congruent than incongruent trials (*t*_25_=2.92, *p*=.007, *d*=.573; 0.29 to 0.42 s, cluster-corrected *p*=.019).

By contrast, the absolute distance between the nontarget item in memory and the random probe orientation (nontarget-probe distance) could be decoded following probe onset on incongruent trials, but not on congruent trials (congruent: *t*_25_=1.59, *p*=.062, *d*=.313; incongruent: *t*_25_=2.92, *p*=.004, *d*=.572; 0.28 to 0.44 s, cluster-corrected *p*=.006). However, there was no significant difference between conditions (*p*=.212; Figure 5D). On congruent trials, decoding quality was significantly greater for the target-probe distance than for the nontarget-probe distance (*t*_25_=2.68, *p*=.006, *d=*.526), while the opposite was true on incongruent trials, although this trend did not reach significance (*t*_25_=-1.53, *p*=.067, *d*=-.300).

If the cued item is incorrectly used as the template for comparison with the probe on incongruent trials, we might expect the neural pattern coding for the target-probe distance on congruent trials and the neural pattern coding for the nontarget-probe distance on incongruent trials to share overlapping representational formats. If so, a decoder trained on target-probe distance values on congruent trials should be able to decode the nontarget-probe distance on incongruent trials, which is what we found (*t*_25_=3.21, *p*=.002, *d*=.630; 0.20 to 0.38 s, cluster-corrected *p*=.001; Figure 5E). That was not the case on congruent trials, where the nontarget had not been cued during the delay (*t*_25_=1.58, *p*=.063, *d*=.310). However, nontarget-probe decoding quality was not significantly greater on incongruent than congruent trials (*t*_25_=-1.17, *p*=.126, *d*=-.230).

In summary, following probe onset, the absolute distance between the probe and the *cued* memory item could be decoded regardless of whether the cued item is the target for orientation recall or not. The comparison signal with respect to the *uncued* item, on the other hand, could not be decoded immediately following probe onset, even when it was the target for orientation recall. These results indicate that the cued item may be prioritised for comparison with the probe at recall.

### Representational overlap between cued item target-probe distance and nontarget-probe distance interferes with precision of orientation recall

We would expect the extent to which the nontarget is mistakenly used as a template for comparison with the probe to interfere with behavioural performance. If so, greater nontarget-probe distance decoding quality (when trained on congruent trials with *target*-probe distance labels) should predict worse orientation recall performance. This was the case on incongruent trials, where there was a positive trial-wise relationship between decoding quality and absolute error (βs>0; *t*_25_= 2.36, *p*=.013, *d*=.464). There was no evidence of a relationship on congruent trials (*p*=.90), where nontarget-probe decoding quality was not significant. β weights were significantly larger on incongruent than congruent trials (*t*_25_=-2.18, *p*=.039, *d*=-.427). There was no evidence of a relationship with RT on congruent (*p*=.36) or incongruent (*p*=.63) trials.

These results were only found after training the decoder on the target-probe distance, indicating that interference was driven by erroneously treating the cued non-target as a target. By comparison, when the decoder was trained on the non-target probe distance, there was no significant relationship with error (congruent: *p*=.87; incongruent: *p*=.73) or RT (congruent: *p*=.39; incongruent: *p*=.37)

## Discussion

Attention can be allocated to select and prioritise task-relevant objects or features in WM for future action. Here we asked whether attentional selection of a single object-feature during the delay enhances processing of other features belonging to the same item. We showed behavioural and neural evidence that attentional selection spreads between an object’s features in WM. Exploratory analyses further revealed that attentional selection during the WM delay prioritises the selected object for subsequent decision-making.

Behaviourally, selection of the cued colour for the incidental task was associated with faster and more precise recall of the same item’s orientation, relative to the orientation of another memory item. This is consistent with studies showing benefits of non-predictive retro-cues, where merely ‘refreshing’ items in WM is associated with improvements in memory performance (Souza, Rerko, & Oberauer, 2015; Yi, Turk-Browne, Chun, & Johnson, 2008; Zokaei, Manohar, et al., 2014; Zokaei, Ning, et al., 2014). Here we show that this cueing effect spreads to other features of a cued object. The incidental task did not demand retrieval of any other features than the cued colour, in contrast to previous incidental cueing studies (Zokaei et al., 2014; 2014b), ruling out explicit retrieval of location as the source of the congruency effect. However, it is possible the behavioural congruency effect could occur in the absence of attentional spread to other features, if the cued colour became more accessible in memory, making it a better cue for subsequent retrieval. To directly assess the effect of colour selection on processing of other features in WM during the maintenance delay, we next looked at neural and eye-tracking signatures of feature-based and spatial attention.

When WM content is prioritised via task-relevant predictive retro-cues, it is represented more strongly in neural activity patterns compared to unattended content (LaRocque et al., 2013; Lewis-Peacock et al., 2012; Sprague et al., 2016; Wolff et al., 2017). Here, even though incidental cues were not predictive or relevant to the memory task, we found that the neural representation for the cued item’s orientation as decoded from EEG voltages was enhanced following the colour cue, while the orientation of the uncued item could not be decoded. An exploratory spatiotemporal decoding analysis revealed greater decoding quality for the cued item’s orientation relative to the orientation of the uncued item. This was the case even though both orientations in memory were equally relevant to the task. These results build a case for obligatory spread of attention between an object’s features in WM. Selecting a feature of a WM object enhances processing of other features belonging to the cued item, even when enhancing them is irrelevant to the memory task.

The observed spread of attention between object-features in WM can be contrasted with the alternative hypothesis that it is possible to selectively attend to a single object feature of a multi-feature object maintained in WM. Several studies have shown that WM selectively encodes and represents object-features that are relevant to the current task (Bocincova & Johnson, 2019; Serences et al., 2009; Woodman & Vogel, 2008). Behavioural benefits afforded by feature-based retro-cues further support feature-based selection in WM (Hajonides et al., 2020; Niklaus et al., 2017). While the abovementioned studies initially appear at odds with the current findings, 100% valid cues may allow irrelevant features to be dropped from memory, thereby freeing up WM resources to process relevant features (Souza & Oberauer, 2016). In the present experiment, the non-predictive cue did not allow removal of any features. The WM object may therefore constitute a relevant “bundle” of connected features that are enhanced whenever a single feature of that bundle is selected for processing (Brady, Konkle, & Alvarez, 2011). This is consistent with theoretical models of attention in WM that retrieve features through auto-associative pattern completion (Lansner et al., 2013; Manohar et al., 2019).

Contralateral alpha suppression and gaze bias are thought to track the focus of spatial attention in WM (Myers, Walther, et al., 2015; van Ede et al., 2019; Wallis, Stokes, Cousijn, Woolrich, & Nobre, 2015). In the present study, a nonpredictive, centrally presented colour cue was associated with alpha lateralization and gaze bias relative to the cued memory item’s original position, even though memory for location was not demanded by the task. These indicate a shift of covert attention to the item’s previous location, possibly as part of the process of selecting the item in memory. Spatial attention may be involved in selection of non-spatial features from WM, even when location is irrelevant (Poch, Capilla, Hinojosa, & Campo, 2017; Van Ede et al., 2020; van Ede et al., 2019).

Contralateral alpha suppression and gaze bias both predicted orientation recall performance, consistent with a functional role in supporting WM for non-spatial features (Cai, Sheldon, Yu, & Postle, 2019). However, the two measures did not correlate at the trial-wise or participant-wise level and showed distinct relationships with behaviour. Alpha suppression contralateral to the cued item’s memorized position predicted slowed orientation recall on incongruent trials. Alpha oscillations may play a role in modulating the relative accessibility of the memory items, perhaps by gating neural information flow between lateralized portions of sensory cortex coding for each item and frontal areas representing current goals (Bonnefond & Jensen, 2012; de Vries, van Driel, Karacaoglu, & Olivers, 2018; Jensen & Mazaheri, 2010; Lara & Wallis, 2014). Gaze bias was associated with greater recall error on incongruent trials, suggesting directional eye-movements toward the cued item impaired memory quality for the other item. This contrasts with previous studies where gaze bias following valid retro-cues mainly predicted reaction times (Van Ede et al., 2020; van Ede et al., 2019). Directional gaze bias during WM is thought to reflect oculomotor involvement in spatial attention “spilling over” into eye-movements, but their function and relationship to neural signatures of spatial attention remain poorly understood (Liu, Nobre, & Ede, 2021). Future studies should further disentangle the relationship between these measures of spatial attention and the ways they support WM for non-spatial features.

Given the neural reinstatement of the cued item’s orientation during the delay, it is unlikely that spatial attention alone explains the behavioural effects of incidental cueing on orientation recall. Nevertheless, the observed spread of attention between non-spatial features (colour and orientation) could be mediated through spatial attention toward their common location, consistent with a special status of location in selecting and binding features in WM (Pertzov & Husain, 2014; Schneegans & Bays, 2017; Treisman & Zhang, 2006). Even though location was irrelevant, it was still a distinguishing feature in our task. It will be interesting to establish whether spatial attention is similarly engaged when memory items share overlapping spatial positions where location information may interfere with performance.

Analyses of neural processing of the probe at recall showed that a comparison signal was automatically computed for the cued object in WM, boosting orientation recall when the cued item was also the target (congruent), but interfering with retrieval when it was not (incongruent). Recent theoretical perspectives predict that attentional selection may prioritise selected information for interaction with new sensory input (Heuer et al., 2020; Myers, Stokes, & Nobre, 2017; Olivers et al., 2011; Olivers & Roelfsema, 2020). Filtering new sensory input through a neural pattern coding for the prioritised memory item (i.e., a matched-filter) may serve as an efficient mechanism for computing the relevant decision signal––in this case, the angular distance of the mnemonic template relative to the probe (Hayden & Gallant, 2013; Muhle-Karbe, Myers, & Stokes, 2021; Myers, Rohenkohl, et al., 2015; Sugase-Miyamoto, Liu, Wiener, Optican, & Richmond, 2008).

The cued item may be maintained in a functionally active state, that influences ongoing processing, by virtue of its recent use in decision-making for the incidental task (Heuer et al., 2020; Myers et al., 2017; Olivers & Roelfsema, 2020; Stokes, Muhle-Karbe, & Myers, 2020). Such a state could be established through strengthened functional connectivity in the network coding for the cued item (Manohar et al., 2019; Oberauer & Lin, 2017; Olivers & Roelfsema, 2020; Rerko & Oberauer, 2013) and enhanced links to neural areas responsible for configuration of task-sets, motor planning and decision-making (Myers et al., 2017; Olivers & Roelfsema, 2020). This may facilitate faster and more efficient comparison between the probe and the cued item, providing a potential mechanism for the congruency effect observed in this study.

In conclusion, attentional selection of a single feature during the delay selectively enhanced processing of other features belonging to the same object. The results show that well-known object-based attention mechanisms exist for internal attention. Attentional selection may prioritise a WM object for comparison with new sensory input, even when the selected object has no special relevance to subsequent behaviour, providing a potential mechanism for non-predictive cueing benefits in WM.

## Data availability statement

Data and custom code will be made available on publication via a link to an open access repository and can be made available to reviewers if desired.

## Acknowledgements

This research was funded by a Biotechnology and Biological Sciences Research Council grant (BB/M010732/1) and James S. McDonnell Foundation Scholar Award (220020405) to Mark G. Stokes and by the NIHR Oxford Health Biomedical Research Centre. Frida A.B. Printzlau was funded by a Biotechnology and Biosciences Research Council studentship (BB/M011224/1) and the Oxford Interdisciplinary Biosciences Doctoral Training Partnership. Nicholas E. Myers was funded by the Wellcome Trust (grant 201409Z/16/Z) and University College Oxford. Sanjay G. Manohar was funded by the MRC clinician scientist fellowship (MR/P00878X) and Leverhulme research grant (RPG-2018-310).

## Citation gender diversity statement

We assessed the gender balance of papers referenced in this article using the Gender Citation Balance Index Tool (https://postlab.psych.wisc.edu/gcbialyzer/). Of the 69 papers with Crossref DOIs, 14 could not be categorised due to missing name data. Of the 56 papers that were successfully categorised, 60.7% had a male first author and male last author, 21.4% had a female first author and a male last author, 16.1% had a male first author and a female last author, and 1.8% had a female first author and a female last author.

## Author contribution

Frida A.B. Printzlau: Conceptualization, software, investigation, formal analysis, writing - original draft. Nick E. Myers: Conceptualization, formal analysis, resources, writing -review & editing, supervision. Sanjay G. Manohar: Conceptualization, writing -review & editing, supervision. Mark G. Stokes: Conceptualization, resources, writing -review & editing, supervision.

